# Damaging *RBM20* E-rich domain variants are not rescued by gene replacement

**DOI:** 10.64898/2026.07.13.738343

**Authors:** Fang (Flora) Bai, Kaiser Chua, Daniel Li, Anna Kirillova, Sunil Yadav, David Staudt, Yuta Yamamoto, Hannah N. De Jong, Linda H. Müller, Kai Fenzl, Non Rungaramsin, Yong Huang, Chloe Reuter, Steven Zhang, Vikki Krysov, Joyce Njoroge, Brendan J. Floyd, Neal K. Lakdawala, Cynthia A. James, Douglas Cannie, Perry Elliott, Luisa Mestroni, Anjali Owens, Marco Merlo, Andrew Krahn, Chad Mao, Diane Fatkin, Vasanth Vedantham, Anwar Chahal, Adam Helms, Joseph C. Wu, Calum MacRae, Michael Gotthardt, Dan Roden, Fritz Roth, Euan Ashley, Benjamin Meder, Brett Kroncke, Daniel Tabet, Atina Cote, Lars M. Steinmetz, Andrew Glazer, Mark Mercola, Victoria N. Parikh

## Abstract

The promise of precision therapeutics in genetic cardiomyopathies relies on linking specific therapies to variant mechanisms. Missense variants in the cardiac splice regulator RBM20 cause a highly penetrant and arrhythmogenic dilated cardiomyopathy. Disease-causing variants in RBM20’s arginine-serine rich (RS) domain act via formation of toxic gain of function cytoplasmic granules, but this is not true for a small number of clinically adjudicated pathogenic variants in its glutamate(E)-rich domain. To better define the effects of E-rich domain variants, we developed a scalable screen based on induced pluripotent stem cell (iPSC) cardiomyocyte differentiation that identified several additional damaging variants. Several of these reduced RBM20 protein abundance and stability. We therefore hypothesized that, unlike RS domain variants, these E-rich variants might be rescued by RBM20 overexpression. To test this hypothesis, we generated induced pluripotent stem cells (iPSCs) from a patient with a pathogenic E-rich domain variant (p.E913K), and confirmed reduced RBM20 protein expression in these *RBM20*^+/p.E913K^ cells after differentiation to iPSC-derived cardiomyocytes (iPSC-CM, vs. engineered isogenic *RBM20*^+/+^). These iPSC-CMs also displayed aberrant transcriptional splicing, reduced contractility, increased calcium-induced calcium release, and nuclear localization of RBM20 protein, often with more than the two expected *RBM20-*centric splice factories. AAV-based overexpression of RBM20 reversed some, but not all of the mis-splicing events identified in *RBM20*^+/p.E913K^ iPSC-CMs, and did not improve their abnormal contractility, calcium handling or supernumerary RBM20 nuclear granules. In summary, our data indicate that pathogenic E-rich domain variants reduce RBM20 protein abundance, but that their mechanism is unlikely to be explained by haploinsufficiency alone.

## Introduction

In the genetics of human cardiomyopathy, many causative variants are thought to lead to disease via loss of protein function, suggesting that they may be treated by increasing the protein abundance of the affected gene. There are several notable exceptions, in some of which different genetic variants converge on similar disease phenotypes via different mechanisms. Further, reduced protein abundance is not always the sole mechanism of disease. The cardiac splicing regulator, RNA binding motif 20 (RBM20) represents one example of this. RBM20 regulates alternative splicing of key structural and calcium-handling genes, thereby controlling cardiomyocyte maturation and function. RBM20 localizes to the nucleus of cardiomyocytes, where it concentrates in discrete “splice factories” associated with active titin RNA processing.(3, 4) Missense variants in RBM20 cause a highly penetrant and arrhythmogenic dilated cardiomyopathy. Variants in RBM20’s arginine-serine rich (RS) domain cause disease via mislocalization to the cytoplasm, inducing abnormal splicing of critical sarcomeric and calcium handling machinery(5). Outside the nucleus, mutated RBM20 forms cytoplasmic ribonucleoprotein (RNP) granules (6), which have been identified as a major driver of cardiomyopathy independent of splice alterations (7). However, the disease mechanism of variants in another major *RBM20* pathogenic hotspot, the glutamate(E)-rich domain (8), is not well understood. One prior report showed that a pathogenic variant in the E-rich domain (p.E913K) is associated with reduced RBM20 protein abundance (9), suggesting that variants in this region may cause disease via haploinsufficiency rather than a toxic gain-of-function mechanism.

To assess the mechanism of disease causality in the E-rich domain, we developed a scalable multiplexed assay of variant effects (MAVE) to evaluate hundreds of variants in the *RBM20* RS and E-rich hotspots. The assay deployed Cas9-derived base editors in iPSCs, followed by differentiation to iPSC-CMs. Based on prior reports of the role of RBM20 in cardiogenesis and cardiomyocyte differentiation (2, 3), we then compared the prevalence of each variant in the iPSC vs. iPSC-CM populations to define their effects on iPSC-CM differentiation. We then examined the effect of damaging E-rich variants on RBM20 protein abundance and stability. Lastly, we assessed the therapeutic potential of AAV-based RBM20 gene replacement in an iPSC-CM derived from a patient with a known pathogenic E-rich domain variant.

## Results

### Endogenous saturation base editing of RBM20 to scalably assay variant effects on iPSC-CM differentiation

In genes like *RBM20,* where the mechanism of disease is hypothesized to differ between variant hotspots, selecting an assay that captures a shared downstream functional consequence allows an expanded view of disruptive variants. Pathogenic variants in RBM20 are known to disrupt cardiogenesis (2, 3) and specifically, induced pluripotent stem cell (iPSC) differentiation to cardiomyocyte (iPSC-CM) fate (1). To establish and validate this assay on a region of RBM20 inclusive of several pathogenic variants, we first transiently expressed highly active cytosine (10, 11) and adenine base editors (12) with a saturation library of guide RNAs (gRNAs) targeted to the RS domain hotspot in iPSCs. iPSCs were enriched for those expressing a base editor via fluorescence activated cell sorting (FACS) and expanded, and the base-edited regions were sequenced with high fidelity polymerase-based enrichment (**Figure 1A)**. Across five biological replicates, 83% of possible single nucleotide variants were observed at a frequency higher than sequencing error (1e-4, **Figure 1B**). iPSCs were then differentiated to iPSC-CMs using standard reprogramming (13). Beating iPSC-CMs were then fixed, stained and sorted to collect the α-actinin2-positive (iPSC-CM fate) population for high-throughput sequencing. Variants were quantified using a custom pipeline and an enrichment score was calculated based on the minor allele frequency of each variant in the iPSC population vs the iPSC-CM population. Because a significant minority of reads showed bystander edits (**Supplemental Figure 1A**), a conservative approach was taken to correct for the effects of bystander edits by recalculating each variant’s enrichment after eliminating reads containing each potential bystander edit and then selecting the enrichment value closest to 1 (additional detail in **Methods and Supplemental Figure 1B**). Corrected estimates of each single nucleotide variant effect were correlated with edits estimated from reads containing only a single variant (*R^2^*= 0.4 (p=2.3e-127, **Figure 1C**), and variant effects were well-correlated between corrected variant enrichments and corresponding enrichments of multinucleotide variants with the same protein (amino acid) consequence (and *R^2^*= 0.7 (p=7.2e-8), **Figure 1D**).

**Figure 1.**
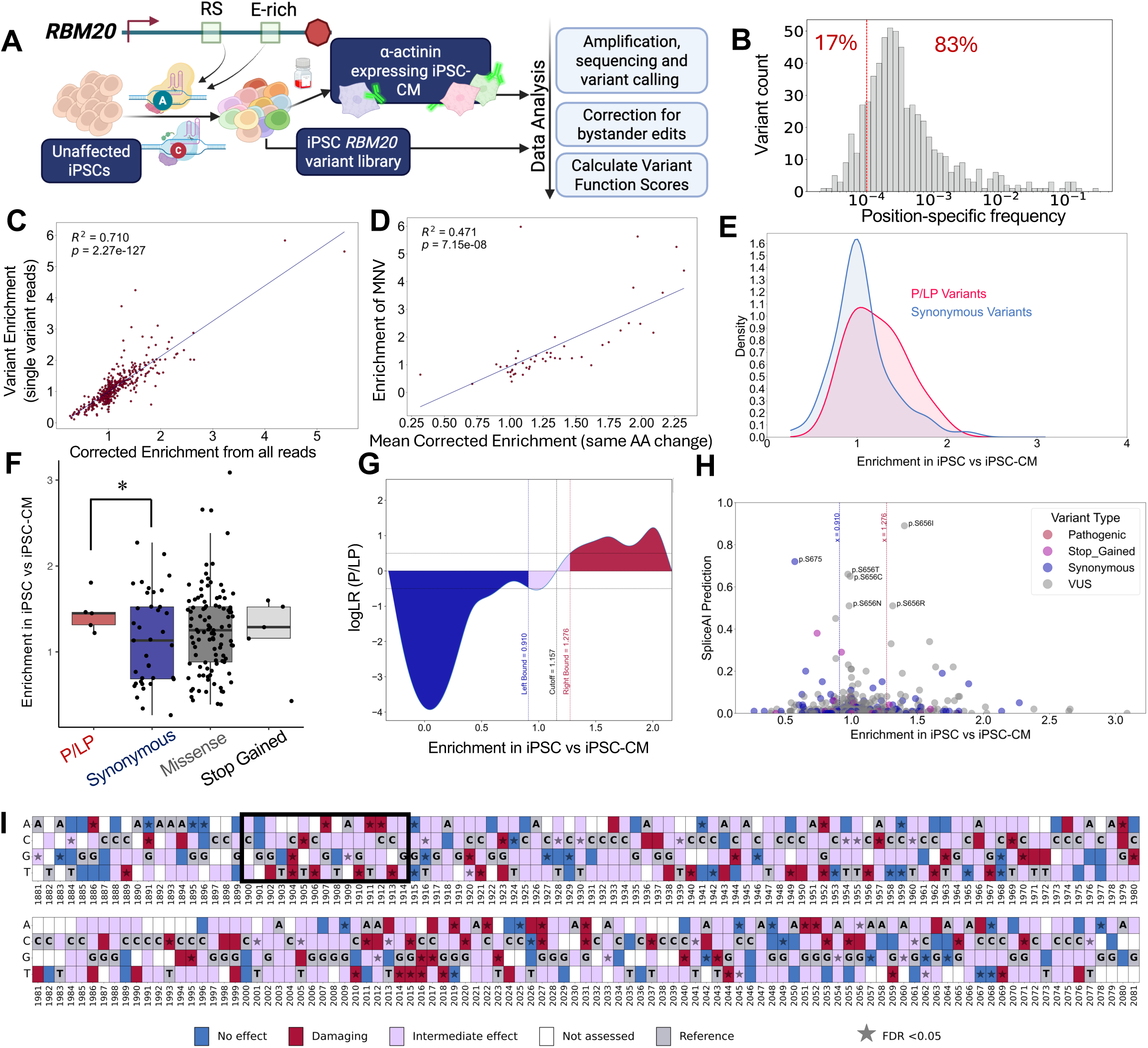
Scaled adjudication of the effects of RBM20 RS domain variants on iPSC-CM differentiation. (A) Schematic of iPSC differentiation assay. (B) iPSC RS domain library displays diversity of variants at frequencies above expected sequencing error (depicted as red dotted line). (C) Corrected enrichment values are correlated with enrichments calculated from single-variant reads only (R^2^=0.71, p=2.3e-127). (D) Corrected enrichment values are correlated with enrichments calculated from reads encoding multi-nucleotide variants (MNVs) with the same amino acid substitution R^2^=0.47, p=7.2e-8) (E) Right shift in pathogenic/likely pathogenic (P/LP) variants (red) enrichment compared to synonymous variants (blue). (F) Among high-confidence enrichment scores (FDR <0.05), P/LP variants have significantly higher mean enrichment scores compared to synonymous variants (1.45±0.22 vs. 1.15±0.53, p=0.04) where as highly predicted stopgain variants are not differentially enriched compared to synonymous variants (1.19±0.42 vs. 1.15±0.53, p=0.82). (G) Log(likelihood ratio) (LLR) of finding a P/LP versus synonymous variant at a given enrichment score with no effect, intermediate and damaging enrichment score cut offs displayed. (H) Splice-AI splice effect predictions of each variant in the RS domain versus effect shows no damaging synonymous variants are predicted to be explained by splice effect. (I) Variant effect map of the RS domain hotspot and surrounding regions. Red = predicted damaging. Blue = predicted no effect, Lavender = intermediate effect. White = frequency too low to adjudicate. Grey = reference sequence. Star = FDR <0.05.

### iPSC-differentiation assay identifies known cardiomyopathic RBM20 RS domain variants as damaging

The distribution of enrichments for known pathogenic and likely pathogenic (P/LP) variants was right-shifted compared to that of synonymous variants in the RS domain (**Figure 1E**). Among high-confidence variant effects (FDR <0.05), P/LP variants had significantly higher enrichment in the iPSC population than synonymous variants (p=0.04), whereas stopgain variants did not (p=0.82, **Figure 1F, Supplemental File 1**).

Log likelihood ratios of identifying a pathogenic variant at a given enrichment value were used to identify cutoffs for damaging variants, intermediate variants, and variants with no effect on iPSC-CM fate differentiation (**Figure 1G**). Given the location of the RS domain at the intron 8-exon 9 junction, we examined the predicted splice effect of each variant. SpliceAI showed a high predicted effect on splicing for a single missense variant in the predicted damaging effect range (p.S656I), though none of the predicted damaging synonymous variants adjudicated in this region displayed high predicted splice effects **(Figure 1H)**. Taken together, these data reflect potential relative tolerance to *RBM20* truncating variants vs. gain of function pathogenic missense variants as we have recently reported in a registry of dilated cardiomyopathy patients with *RBM20* variants.(14) Single nucleotide variant effect estimates across the RS domain hotspot are displayed (**Figure 1I)**, and are also provided as estimates of amino acid substitution effect (**Supplemental Figure 2**).

### Extension of RBM20 variant effects assay to E-rich hotspot and clinical correlation with effect predictions

We next extended this assay to the E-rich domain by performing saturation mutagenesis of a 200 bp section of this domain centered around a 40bp disease-associated hotspot we previously identified.(8) 49% of possible variants in the E-rich domain hotspot were observed at a frequency higher than sequencing error (1e-4). The majority of variants identified as damaging in the combined RS domain and E-rich domain hotspots were rare in UK Biobank (UKB), with the exception of p.Asn888Asp, which is prevalent in the Finnish European population in gnomAD (2%), and non-Finnish Europeans (0.3%) suggesting the potential for founder variant status with very low disease penetrance or a false positive from the screen (**Figure 2B**).(15) Excluding this variant, participants in UK biobank carrying a predicted damaging variant were more likely to have a cardiomyopathy diagnosis during their lifetime than those with a variant predicted to have no effect (damaging variants 4.3% likelihood, no-effect variants 0.9% likelihood, odds ratio (OR) 5.2[1.1,20.9],p=0.02). We have previously reported a higher burden of major ventricular arrhythmia and evidence of higher penetrance of RBM20 cardiomyopathy compared to other inherited dilated cardiomyopathies.(8) We therefore asked whether patients with predicted damaging and intermediate variants of uncertain significance (VUS) would dilute these clinical characteristics in a registry of patients with known P/LP variants in RBM20.

**Figure 2.**
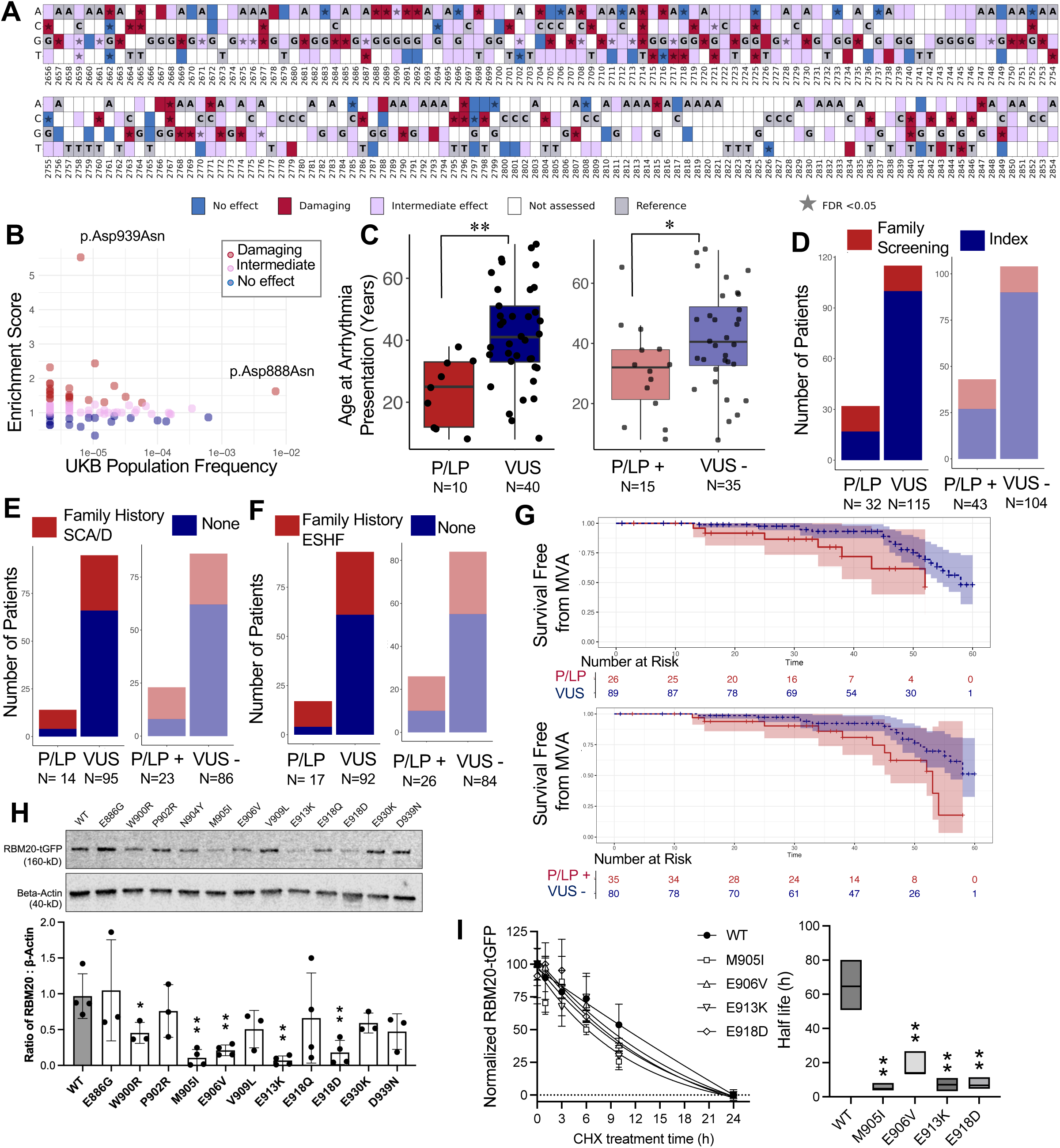
Predicted variant effects in the RS and E-rich domains are associated with RBM20-relevant clinical phenotypes and E-rich domain variants can affect protein stability. (A) Variant effect map of the E-rich domain hotspot and surrounding region. Red = predicted damaging. Blue = predicted no effect, Lavender = intermediate effect. White = frequency too low to adjudicate. Grey = reference sequence. Star = FDR <0.05. (B) UK Biobank minor allele frequencies versus enrichment score for observed missense variants. (C) Age at presentation of arrhythmia is higher in VUS than P/LP in the RBM20 registry, and this remains true after movement of damaging and intermediate VUS into the P/LP category (“P/LP+” and VUS -”). (D-F) Proportion of patients in RBM20 DCM registry who presented for family screening, had a family history of sudden cardiac arrest, or a family history of end stage heart failure in P/LP vs VUS and in P/LP+ vs VUS-. p<0.05 for all comparisons by Fisher Exact test. (H) Log rank comparison of life-time incidence of major ventricular arrhythmia (prior to age 60) in P/LP vs VUS and P/LP+ vs. VUS - (P/LP vs. VUS: p=0.04, HR 0.38[0.15, 0.94]; P/LP+predicted damaging and intermediate vs. VUS -: p=0.03, HR 0.41[0.19,0.88]). (H) RBM20 protein abundance in HeLa cells of highly predicted damaging variants in the E-rich domain reveals reduced abundance compared to reference RBM20 sequence (WT) for several variants. (I) Variants with reduced protein abundance display reduced protein stability in cycloheximide pulse chase experiments. For all panels: *p<0.05, **p<0.01, ***p<0.001, ****p<1e-4.

Indeed, we found that among patients who presented with arrhythmia, those with P/LP variants presented at a younger age vs. those with VUS, and that when predicted damaging or intermediate effect VUS were added to the P/LP group (“P/LP+”), this relationship was unchanged (**Figure 2C**, p=0.001 and 0.03 respectively). Likewise, the proportion of patients presenting for family screening rather than as an index case (the first in their family to have a cardiomyopathy diagnosis), was higher for the P/LP variant patients (OR 0.2 [0.1,0.5], p=9.8e-5). Supporting increased prevalence of adverse family histories associated with P/LP variants, the proportion of patients with family histories of sudden cardiac arrest/death (OR 5.6[1.5,26.5],p=0.005) and endstage heart failure (OR 6.1[1.7,27.8],p=0.003) were also increased among patients with P/LP variants. None of these findings changed by moving patients with damaging or intermediate effect VUS to the P/LP category (**Figures 2D** (OR 0.27[0.1, 0.7],p= 0.003), **2E** (OR 4.8[1.6, 14.8],p=0.001) and **2F** (OR 3.0[1.1, 8.4],p=0.02) respectively). Lastly, moving these patients with predicted damaging and intermediate VUS to the P/LP group delivered consistently decreased odds of early major ventricular arrhythmia (sudden cardiac arrest, ventricular tachycardia or appropriate defibrillator shock before age 60) in the VUS patient population. (**Figure 2G**, VUS vs. P/LP: HR 0.38[0.15, 0.94], p=0.04; VUS - vs PLP+: HR 0.41[0.19,0.88],p=0.03).

### Predicted damaging E-rich domain variants are variably associated with reduced RBM20 protein abundance

Several variants in the E-rich domain were significantly associated with impaired iPSC-CM differentiation (**Figure 2A, Supplemental Figure 2, Supplemental File 1**). The single known disease causing variant in this region, p.E913K, was not classified as damaging by our assay, but this may reflect the specificity of our assay and its focus on developmental variant effects: The mechanistic effect of p.E913K in a diploid context may not have a significant enough effect on iPSC-CM differentiation to be captured as damaging here, but could still cause human disease in a mature cardiomyocyte via loss of function. Based on prior reports of reduced protein abundance of p.E913K, however, we tested the hypothesis that p.E913K and highly predicted damaging E-rich variants from the iPSC-CM differentiation assay demonstrate reduced RBM20 protein abundance.(9) We transfected HeLa cells with a series of RBM20 variant constructs from a C-terminal GFP-fusion *RBM20* construct, and demonstrated that several of these variants, as well as the previously reported p.E913K, displayed reduced expression (**Figure 2H)**. There is no available crystal structure of the disordered E-rich domain, and the predicted structure of RBM20 is highly intrinsically disordered, precluding accurate computational modeling of the effects of these variants on protein stability, however, cycloheximide pulse chase experiments confirmed a reduced half life of the RBM20 variant proteins with reduced overall protein abundance (**Figure 2I**). We further found that co-expression of some of our most highly predicted damaging variants with normal RBM20 resulted in reduced overall RBM20 protein abundance, indicating that some of these variants have a dominant negative effect on the functional copy of RBM20 **(Supplemental Figure 3A).** Taken together, these results suggest that several damaging variants in the E-rich domain cause reduced RBM20 protein abundance via disrupted protein stability, possibly with dominant negative effect, consistent with a potential loss of function mechanism of disease for these E-rich domain variants.

### p.E913K iPSC-CMs display reduced protein, aberrant splicing, abnormal contractility and calcium handling

We generated an iPSC line from a patient with an arrhythmogenic dilated cardiomyopathy in which heterozygous *RBM20* p.E913K was the sole causative variant identified on a broad clinical genetic testing panel for known causes of arrhythmia and cardiomyopathy. We used prime editing to replace this p.E913K variant with the reference sequence (an isogenic +/+ control, **Supplemental Figure 3B).** After differentiation to iPSC-CMs, p.E913K and isogenic +/+ line displayed no difference in *RBM20* RNA expression by qRT-PCR, but p.E913K iPSC-CMs had clearly reduced RBM20 protein expression compared to isogenic +/+ controls (**Figures 3A&B**). RNA-seq of p.E913K/+ iPSC-CMs displayed aberrant splicing of RBM20-target exons after hybrid capture enrichment for known RBM20 splicing targets (**Figure 3C and Supplemental Figure 3C**), key targets of which were confirmed by qRT-PCR (TTN, CAMK2D, and RYR2; **Figure 3D-F**). Based on kinetic image cytometry (16) p.E913K/+ iPSC-CMs displayed reduced contractility as compared to isogenic +/+ iPSC-CMs (**Figure 3G**, p=2.5e-8). Fluorescent calcium transient analysis demonstrated increased amplitude of calcium-induced calcium release in p.E913K/+ iPSC-CMs (p=1e-4), as previously reported in *RBM20* deficient mice (17), as well as longer transient duration (corrected for beating rate, p=0.02, **Figure 3H**). Although these calcium and contractility findings may appear incongruent with one another, they may be explained by expression of an aberrant L-type calcium channel subunit isoform that leads to increased calcium influx, elevated cytosolic calcium, and elevated sarcoplasmic reticulum calcium content.(17) The association of this finding with reduced contractility may be related to the many other differentially spliced transcripts relevant to sarcomeric structure, or long term metabolic stress from hypercontractility and “burn out.” Lastly, we confirmed that RBM20 protein is located within the nucleus of p.E913K/+ iPSC-CMs, but also identified a previously undescribed nuclear distribution in which RBM20 does not localize to two nuclear splice factories seen in +/+ iPSC-CMs, which are expected based on prior observations (3). Rather, in p.E913K/+ iPSC-CMs, cells with two or fewer RBM20-positive nuclear inclusions were noted less often, with as few as 40% of nuclei in a given microscopy field displaying the expected two-speckle architecture **(Figure 3I).**

**Figure 3.**
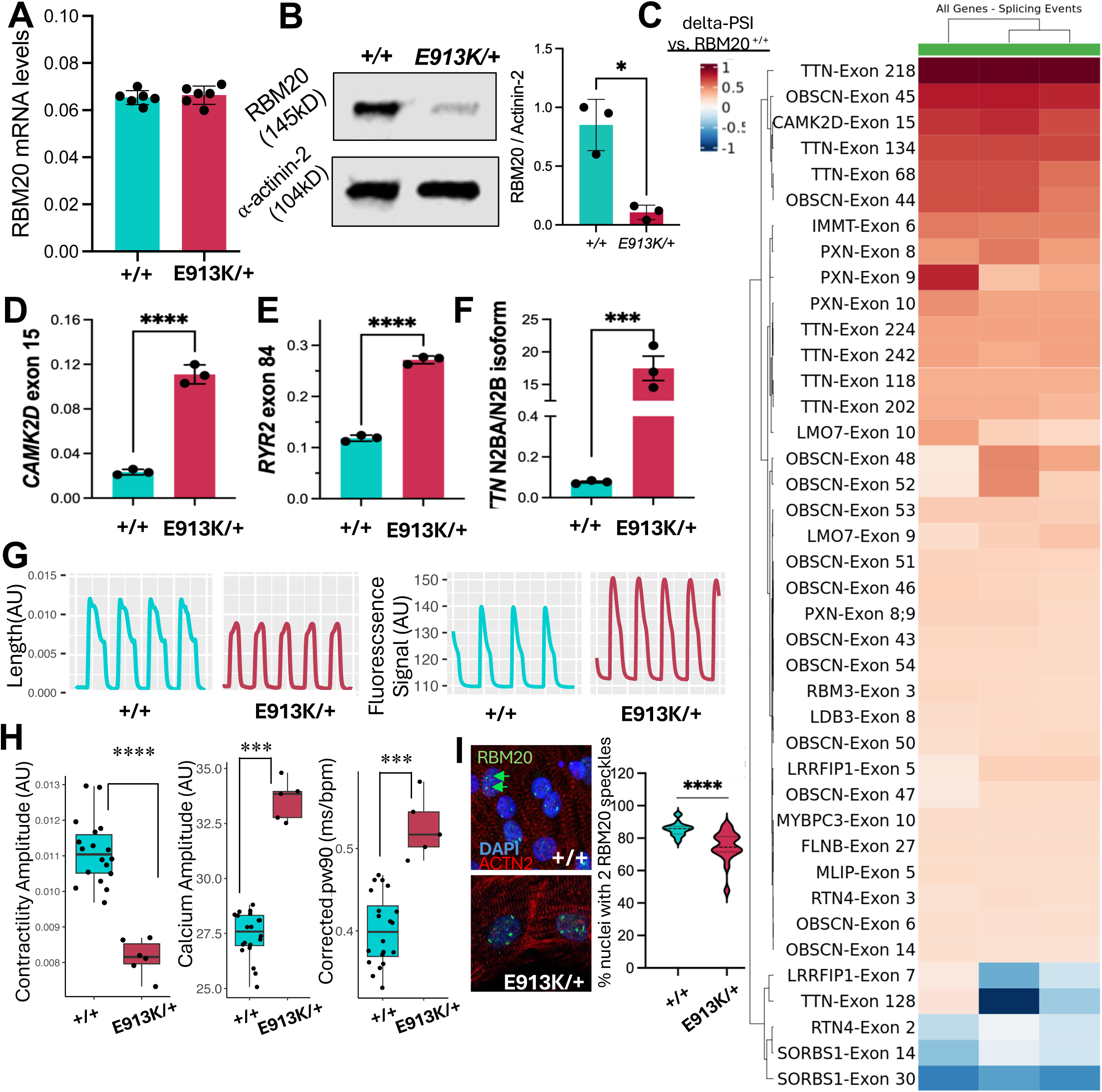
*RBM20* p.E913K/+ iPSC-CMs display reduced RBM20 protein, abnormal splicing, and cardiomyopathic traits. (A) RNA expression (qPCR) is not different between p.E913K/+ iPSC-CM and +/+ iPSC-CM (isogenic control). (B) Reduced RBM20 protein expression in p.E913K/+ iPSC-CM vs +/+ iPSC-CM. (C)Targeted enrichment of RBM20 target transcripts reveals expected aberrant inclusion (red =inclusion, blue = exclusion, FDR <0.05 for all comparisons shown), which are verified by qPCR (D-F). (G) Representative contractility traces from single well position during kinetic image cytometry (Blue = +/+, Red = p.E913K/+). (H) Representative calcium fluorescence traces from single well position during kinetic image cytometry (Blue = +/+, Red = p.E913K/+). (I) p.E913K/+ iPSC-CM display reduced contractility compared to +/+ iPSC-CMs. (J) p.E913K/+ iPSC-CM display increased calcium release amplitude compared to +/+ iPSC-CMs. (K) p.E913K/+ iPSC-CM display increased rate-corrected calcium release width (pw90) compared to +/+ iPSC-CMs. For all panels: *p<0.05, **p<0.01, ***p<0.001, ****p<1e-4.

### RBM20 gene replacement rescues splicing, but not contractility or calcium handling in RBM20 p.E913K iPSC-CM

To test the hypothesis that AAV-mediated RBM20 gene replacement would ameliorate the effects of p.E913K, we used an AAV6-*RBM20* expressing the RBM20 cDNA under the control of a cardiomyocyte-specific *TNNT2* promoter (**Supplemental Figure 3F**). We determined the MOI at which normal nuclear splicing architecture of RBM20 (two nuclear RBM20-positive splice factories) was recapitulated in *RBM20^-/-^* iPSC-CMs engineered using a Cas9 induced deletion of the start codon of *RBM20* **(Supplemental Figure 3D&E).** We then treated p.E913K/+ iPSC-CMs at a range of MOIs (0.5e4-2e4) between days 40 and 60 of differentiation using metabolic maturation media (16). Five days after AAV treatment, robust mRNA and protein expression of RBM20 was observed (**Figure 4A, Supplemental Figure 3G**). Inclusion of *CAMK2D* exon 15, *RYR2* exon 84 and the ratio of TTN NB2A/N2B isoforms were all corrected by AAV treatment (**Figure 4B**). Notably, all were corrected to levels lower than seen in the isogenic corrected (RBM20^+/+^) iPSC-CM line. Contractility and calcium handling, however, remained unchanged, with a minor apparently detrimental effect of empty and GFP-carrying AAV versus lower dose AAV-RBM20 on contractility (p=0.003, p= 0.01 respectively, **Figure 4C).** In addition to three replicates at the 5 day timepoint, this experiment was repeated at 14 days after AAV treatment and showed similar results, though at this timepoint, AAV-GFP did not have a significant effect on contractility (p.E913K/+ with AAV-RBM20 vs AAV-GFP: p=0.36, p.E913K/+ without AAV vs AAV-GFP: p=0.10 **Supplemental Figure 4**). The inflammatory cytokine CCL-1 displayed increased transcript expression in a dose-dependent manner 5 days after treatment, but had resolved after 14 days, whereas neither IL-1B nor IL-6 were increased by AAV treatment at the 5 day time point. At 14 days after treatment, IL-1B showed a modest elevation of 1.5-2X untreated control levels (**Supplemental Figure 4I**).

**Figure 4.**
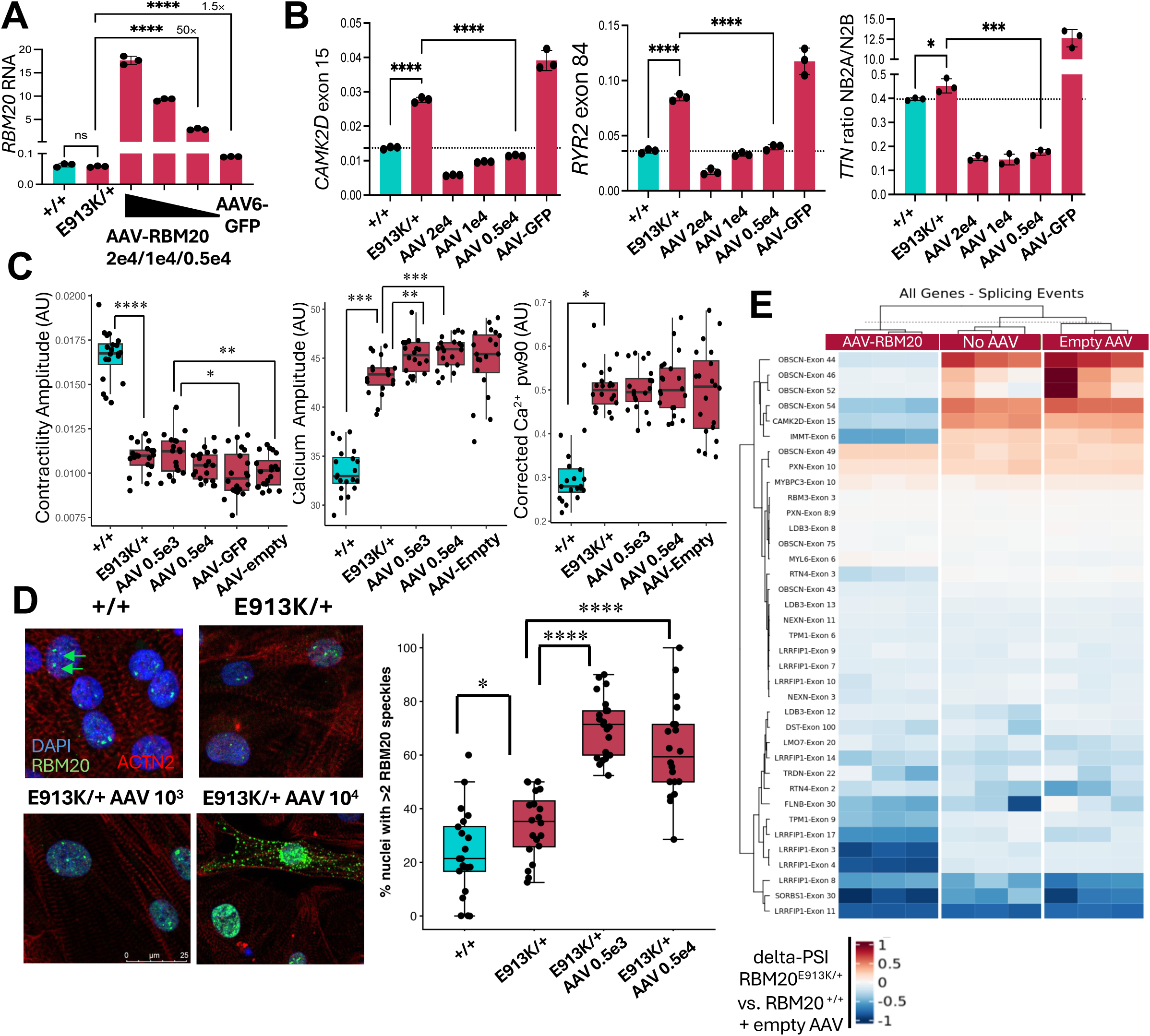
RBM20 gene replacement repairs canonical splicing defects but not contractility or calcium handling in *RBM20* p.E913K/+ iPSC-CMs. **(A)** RBM20 RNA expression in response to AAV dose escalation (qRT-PCR). **(B)** Canonical RBM20 splicing targets by qRT-PCR display suppression of inclusion by AAV-RBM20 in p.E913K/+ iPSC-CM vs. +/+ iPSC-CM. **(C)** Comparison of contractility, calcium transient amplitude and calcium transient width (corrected for beating rate) with and without AAV in p.E913K/+ iPSC-CM vs. +/+ iPSC-CM 5 days after AAV treatment compared to AAV-GFP. **(D)** Multiple nuclear RBM20+ (green) speckles in p.E913K/+ iPSC-CM verses expected 2 RBM20+ splice factories in corrected isogenic iPSC-CM. Number of nuclear speckles per nucleus increases with AAV treatment. The proportion of nuclei with >2 speckles in each genotype is depicted (N=20 per group, ANOVA p<2e-16). **(E)** RNAseq of p.E913K iPSC-CMs with and without AAV treatment (red =inclusion, blue = exclusion). Comparisons depicted are for exons with |PSI| >0.15 and FDR <0.05. **(F)** Co-transfection of wildtype (WT) RBM20 with E-rich variant constructs in HeLa cells. p.M905I and p.E918Q demonstrate a dominant negative effect on WT protein abundance, while p.E913K, p.E906V, p.E918D and p.R636C (RS domain variant) do not. pBS= pBlueScript (empty plasmid). For all panels: *p<0.05, **p<0.01, ***p<0.001, ****p<1e-4.

These data suggest that repair of canonical splicing via AVV-mediated replacement of RBM20 protein expression alone is insufficient to repair cardiomyopathy phenotypes in E-rich domain variant RBM20 iPSC-CMs. Certainly elevated inflammatory markers have been associated with reduced contractility in iPSC-CMs, though these changes occurred within 48-72 hours after treatment with SARS-CoV to or lipopolysaccharide, (18, 19) whereas the elevations we observe in CCL-1 resolve before the 14 day timepoint, and modest elevations in IL1B are delayed and generally modest compared to these prior reports. We therefore hypothesized that an alternate mechanism beyond haploinsufficiency might be responsible for *E913K*-RBM20 cardiomyopathy.

Unlike RS domain variants, which cause damaging cytoplasmic RNP aggregates leading to disease independent of abnormal alternative splicing, p.E913K does not localize to the cytoplasm(9), originally prompting the investigation of the haploinsufficiency mechanism interrogated here. On high resolution fluorescence microscopy, we indeed found no evidence of cytoplasmic RNP granules in p.E913K iPSC-CMs. We did, however, find evidence of increased nuclear RBM20-positive speckles as compared to the two RBM20-TTN directed splicing factories identified in normal human and wildtype pig cardiomyocytes(3, 6) (**Figure 4D)**. AAV-RBM20 treatment did not resolve the supernumerary RBM20-positive speckles, and indeed displayed cytoplasmic distribution of RBM20 at higher doses. Further, although correction of canonical splicing was observed with AAV treatment (**Figure 4B**), global RNA sequencing (RNAseq) of known RBM20 targets revealed no change and at times worsening of non-canonical exon inclusion in targets such as LRRFIP1, TPM1, NEXN, LDB3, and SORBS 1 (**Figure 4E, Supplemental Files 3-5**). This is consistent with persistence of mis-directed splicing by the p.E913K protein despite overexpression of normal RBM20. Taken together, these results indicate a combination of gain of function (misdirected splicing) and/or dominant negative effects for several damaging variants in the RBM20 E-rich domain.

## Discussion

We present a systematic identification of novel damaging variants in the RS and E-rich domain of *RBM20.* We find that many of these E-rich domain variants demonstrate reduced protein abundance and stability, and that while RBM20 gene replacement repairs some canonical splicing deficits in a pathogenic E-rich domain variant iPSC-CM model, it is insufficient to repair critical cardiomyopathy-relevant phenotypes, suggesting an additional gain of function mechanism of E-rich domain variants related to nuclear splicing organization.

The MAVE presented here identified multiple E-rich domain variants to be biologically damaging in an endogenous, heterozygous cardiomyocyte context. This allowed a multi-variant assessment of E-rich domain variant effect on RBM20 abundance and stability, motivating a gene replacement approach to therapy. This application of a variant effect assay in a previously under-studied region of RBM20 rapidly provided evidence for more variants than might otherwise be uncovered in the human population over decades. However, because application of MAVE data in the clinical setting requires significant *a priori* knowledge of variant-disease mechanisms and high numbers of known pathogenic or likely pathogenic variants (20) these variant effect maps of *RBM20* cannot provide strong evidence for clinical variant interpretation at this time, and should not be used in isolation to reclassify variants in a clinical context.

That this map is provided in a heterozygous cardiomyocyte model context rather than a hemizygous episomal context confers substantial human disease-relevance. Specifically, while hemizygous screens yield very clean variant effects with respect to loss or gain of function, they cannot always reveal the implications of loss of variant function in a whole-cell context. For example, here, stop-gain variants were not overall associated with impaired iPSC-CM differentiation, consistent with previously reported relatively small effect of RBM20tvs in the human population (21). Therefore, it is unlikely that loss of function represents a major mechanism of monogenic (isolated) variant effects in human RBM20 cardiomyopathy. This idea is also bolstered by the lack of complete reversal of disease phenotypes after AAV gene replacement in E913K iPSC-CMs. However, the tradeoff of these observations is a limitation of this system: The underlying mechanism of disease for damaging variants is not always clear. Several variants were identified to affect iPSC-CM differentiation that are outside known nuclear localization domains and human variant hotspots. Further, multiple assays are likely needed to fully explore the mechanistic landscape of variant effects in a given gene. The superimposition of the variant effect map presented here with other maps testing different underlying mechanisms of disease may reveal how the novel variants we identify disrupt cardiomyocyte differentiation, as well as variants for which this disruption remains mechanistically unexplained.

That RBM20 gene replacement can correct splicing of many canonical RBM20 target transcripts in p.E913K iPSC-CMs *without* correction of contractility or calcium handling is a surprising result. It is possible that iPSC-CM protein turnover takes longer than 14 days to demonstrate the full effect of splice correction on contractility and calcium, however, the turnover of titin has been reported to be as short as hours or days in embryonic cardiomyocytes (22). Otherwise, the data presented here imply that there is a mechanism of disease other than protein instability haploinsufficiency contributing to E-rich domain variant pathogenicity. In RS-domain variants, reducing RBM20 expression has a negative effect on splicing and a *positive* effect on cardiomyopathy phenotypes in rodents, suggesting a cardiomyopathic effect of mutant cytoplasmic granules themselves.(7) Our data evidence an additional such mechanism for E-rich domain variants. This may be linked to the RBM20-positive intranuclear speckles we observe, which may indicate that p.E913K cannot appropriately nucleate necessary RBM20 splice factor architecture associated with *TTN* and other RBM20-spliced genes.(3) That abnormal splicing of non-canonical exons and disrupted nuclear architecture are observed even after wildtype RBM20 is re-introduced, further evidences a gain of function mechanism by which the p.E913K mutation is independently disrupting these phenotypes. Additional investigation of the composition of these multiple nuclear RBM20-positive speckles as well as the differential binding of RBM20 E-rich domain variant proteins to *TTN* transcripts and other organizers of genomic and transcriptional architecture are an important focus of future research into this putative gain of function mechanism.

Overall, these data support and broaden a growing body of literature that suggests *RBM20* missense variants, including those in the E-rich domain, may not be addressed with gene replacement therapy. While the first reports of *RBM20* loss of function in rodents helped us understand the crucial role of RBM20 in directing splicing of sarcomeric and calcium handling components (5, 17), later research involving human and porcine models has shone a light on the potentially pathogenic role of RNP granules in this disease (3, 4, 6, 7). We have also found that while RBM20 truncating variants are found in hereditary DCM clinics, they are only weakly enriched in cardiomyopathy at a population level, and are associated with milder disease course in the clinic compared to disease-causing missense variants (21, 23). As we move toward therapies for patients with *RBM20* cardiomyopathy, the work we present here creates a platform for extension of novel therapeutics that target gain of function and dominant negative mechanisms of *RBM20* variants in the E-rich domain.

## Acknowledgements

This work was supported by NIH R01 HL168059 and K08 HL143185 (VP), the multicenter CardioVar NIH R01 HL164675 (DR, EA, FR, CM, AG, BK and VP), a LeDucq transatlantic networks of excellent grant: CArdiac Splicing as a Therapeutic Target (CASTT) (EA, LM, BM, MG and VP), NIH R01 HL164634 (LM), NIH R01 HL130840 and S10 OD030264 (MM), K08 HL165094 and AHA 23SCEFIA1154505 (DS), EU Horizon NextGen project (AC).

## Materials and Methods

### RBM20 variant iPSC library development

The *RBM20* variant iPSC library was engineered using the cytosine base editor CRISPR-X(11) and adenine base editor ABE8e (12). Guide RNAs were identified using CHOP-CHOP(24), selecting for guides with <2 off-target mismatches in the genome and targeted to the 200 bp genomic regions described (Supplemental File 1). iPSCs were electroporated with a 1:1:1:1 ratio (by weight) of CRISPR-X-BFP fusion plasmid, ABE8e plasmid, pBlueScript plasmid engineered to express the sgRNA library, and empty pBlueScript. A Neon electroporation system was used for electroporation of 1 million iPSCs from an unaffected participant in 100ul buffer at 1100V, for 30 ms and 1 pulse. Electroporated iPSCs were immediately plated to matrigel-coated 24 well plates in mTESR1 with thiazovivin. Cells were sorted by FACS to select BFP+ cells 48 hours after transfection and expanded. iPSC libraries were sequenced at the RBM20 loci of interest (for exon 9 Fwd primer: 5’- GCAGAATGAGTAAAGGCACAGCGATG-3’ Rv Primer: 5’- GGCTCTTTCCGGTAGTAGCCGTCTTC-3’; for exon 11 Fwd primer: 5’-GAGTGGTCCTTATGGCCAAG-3’ Rv Primer: 5’- AGGCCTGTCTCACTTTCAGC-3’) using tagmentation via the Illumina Nextera XT DNA Library Preparation Kit (15032354) and sequenced on an illumina iSeq.

### iPSC-CM differentiation assay

After selecting BFP+ iPSCs by FACS, iPSC libraries were expanded in MTESR and plated at 80% confluency on matrigel-coated 12 well plates. Differentiation was performed with 0.5 uM ChiR and 0.5 uM IWR with lactate selection starting at day 13 as we have previously described.(13) On day 30 of differentiation, cells were treated with 1:1000 Brefeldin A, dissociated, fixed in solution with 4% PFA, permeabilized and stained overnight with a primary antibody to α-actinin 2 at a dilution of 1:1000 (α-actinin 2, *Sigma A7811)*. After incubation with secondary antibody (goat-anti-mouse, Alexa488, *Invitrogen A11029*), cells were washed and FACS sorted for GFP fluorescence. α-actinin 2 positive cells underwent DNA extraction (Quick Extract, BioSearch Technologies), and cells were genotyped using the same protocol as the iPSC library above.

### iPSC-CM library evaluation and differentiation assay analytic pipeline

A custom pipeline was written for analysis and is publicly available (https://github.com/AshleyLab/vesper).

For variant counts, after QC, fastqs were aligned with BWA, followed by extraction of variant counts in each pool from mpileup files. Variant counts were then reformatted to count unique instances of each variant. Minimap-2 was used as a caller in combination with custom script to identify multi-variant reads by the unique variants they contained. For analysis, five independent replicates of the assay were pooled for the RS domain hotspot and 4 independent replicates for the E-rich domain.

Enrichment analysis was first performed on single nucleotide variants called from the BWA alignment based on the variant frequency in each pool (variant count/number of reads covering that variant position). Variants with a mean frequency less than sequencing error between pools were not analysed. Each variant’s frequency in the iPSC population was then divided by its frequency in the α-actinin 2 positive iPSC-CM population to calculate its enrichment in the immature (iPSC) population. Therefore a higher enrichment score (>1) reflects poorer differentiation capacity.

Given the number of reads with multiple variants, corrected enrichments were calculated as follows (Supplemental Figure 2):

1. All reads with a given variant were counted in both pools (iPSC and iPSC-CM)
2. Reads containing that variant paired with other variants were counted individually per variant pair.
3. Reads containing a given pair were sequentially dropped from the enrichment calculation for that variant to determine the enrichment of the variant of interest independent of each variant with which it was observed to be paired.
4. Of the resultant enrichment values, the value closest to 1 was taken as the “true” enrichment value, thus eliminating potential polluting effects of co-traveling damaging variants on variants with no or intermediate effect.

### UK Biobank and Clinical RBM20 DCM registry and variant adjudication

For comparisons within UK Biobank (UKB) the cohort of 502,132 participants with genome data was accessed using the UK Biobank Browser.(25) For cardiomyopathy diagnosis prevalence, the ICD-10 code I42 was used.

For the RBM20 DCM Registry, patients with a variant in *RBM20* identified on clinical genetic testing were identified at international familial cardiomyopathy centers: Barts Heart Centre (London, UK), Brigham and Women’s Hospital (Massachusetts, USA), Children’s Healthcare of Atlanta (Georgia, USA), University of Colorado Anschutz Medical Campus (Colorado, USA), Heidelberg University (Germany), Johns Hopkins University (Maryland, USA), University of Pennsylvania (Pennsylvania, USA), Royal Brompton Hospital (London, UK), Stanford University (California, USA), University of Trieste (Italy), University of British Columbia (Canada), Victor Chang Cardiac Research Institute (Australia), University of California, San Francisco (California, USA), Wellspan Health (Pennsylvania, USA), University of Michigan (Michigan, USA). Clinical data were collected retrospectively from real-world prospective care at these centers in case note abstraction forms. Dates were not provided but recorded as ages in years. Data were collected, stored, and shared in a deidentified manner in accordance with the institutional review board at each contributing center as named above.

Variant pathogenicity was adjudicated centrally by a licensed clinical genetic counselor and a board certified medical geneticist (**See Supplemental File**). Each variant was classified into one of three categories based on American College of Medical Genetics criteria with modifications for DCM: pathogenic/likely pathogenic variants (P/LP), variants of uncertain significance (VUS), or truncating variants.

### Generation of patient-derived p.E913K/+ iPSCs and engineering of variant-corrected (+/+) isogenic iPSC line

The pathogenic RBM20 E913K variant (c.2737G>A) in the DCM patient-derived hiPSC line SCVI-2292 was corrected using the adenine base editor ABE8e [1]. A single-guide RNA (sgRNA) targeting the E913K locus was designed using CHOPCHOP (https://chopchop.rc.fas.harvard.edu), and corresponding oligonucleotides are listed in Table 1. Annealed sgRNA oligos were cloned into Esp3I-digested pBS to generate the sgRNA expression plasmid pBS-ABE-sgRNA (Table 2). Heterozygous E913K/+ iPSCs were electroporated with a 1:1 mixture of NG-ABE8e-BFP and pBS-ABE-sgRNA using the Invitrogen Neon Transfection System (1100 V, 30 ms, one pulse). Twenty-four hours after electroporation, BFP-positive cells were isolated by fluorescence-activated cell sorting (BD FACSymphony S6) and plated as single cells into 96-well plates in CloneR-supplemented mTeSR medium (STEMCELL Technologies). After ∼10 days of clonal expansion, emerging colonies were manually picked and split; one portion was used for genomic DNA extraction with QuickExtract DNA Extraction Solution (Biosearch Technologies), followed by PCR and Sanger sequencing using primers listed in Table 1.

**Table 1.**
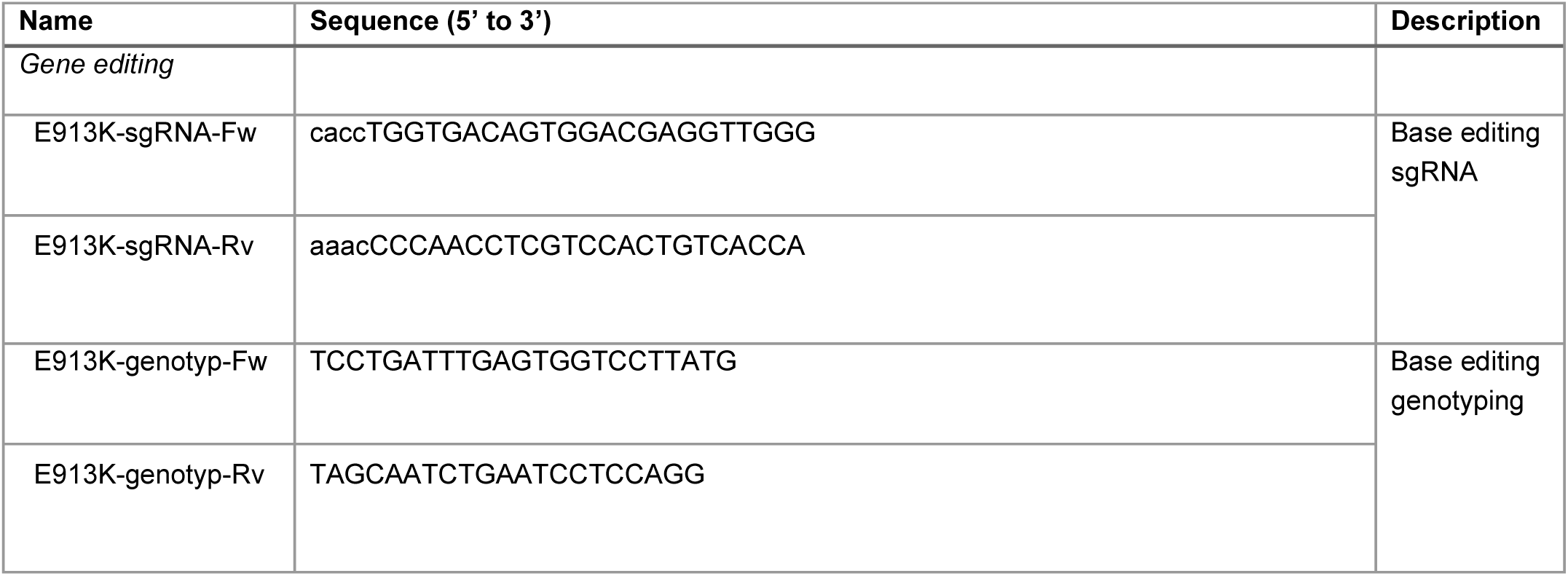
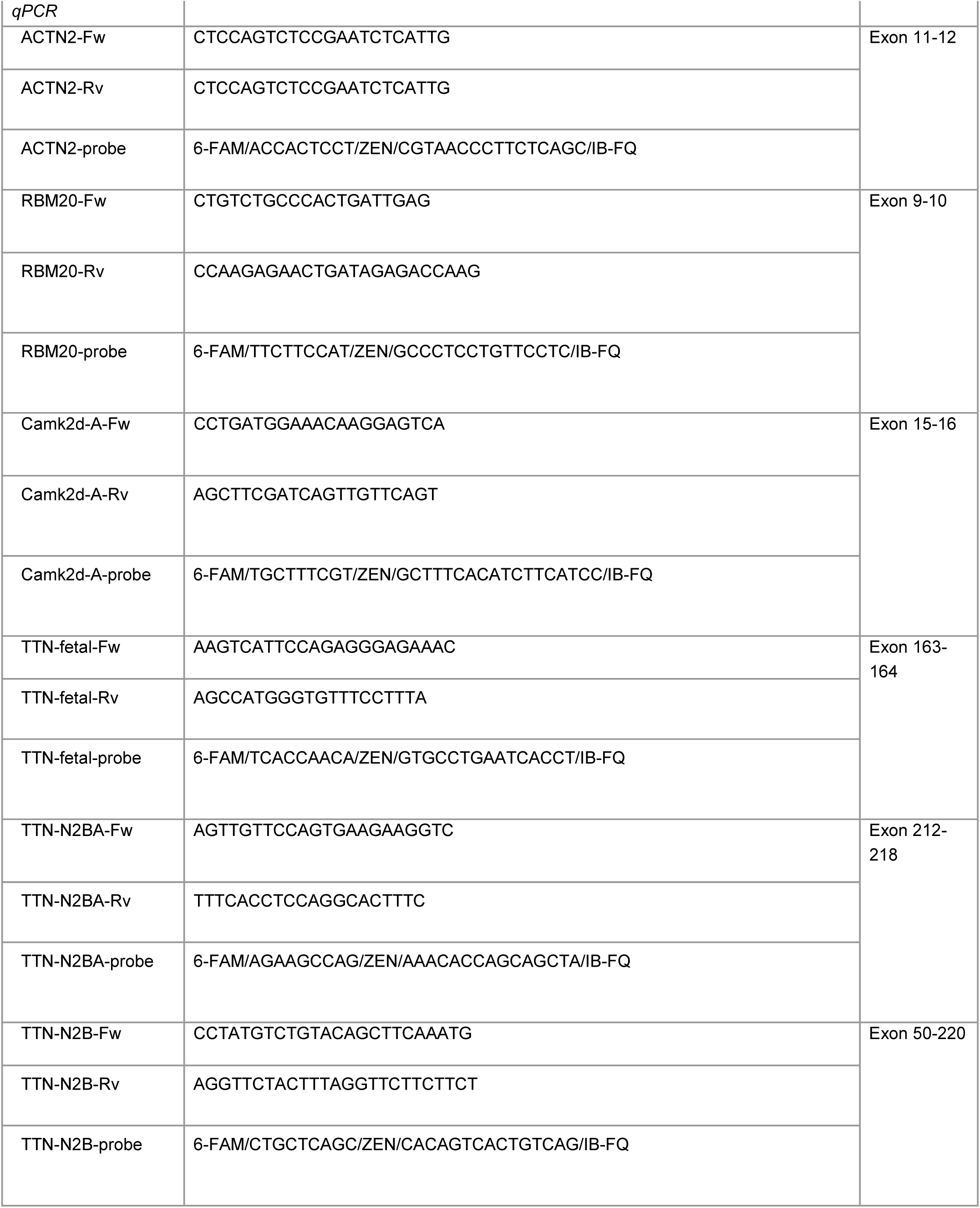
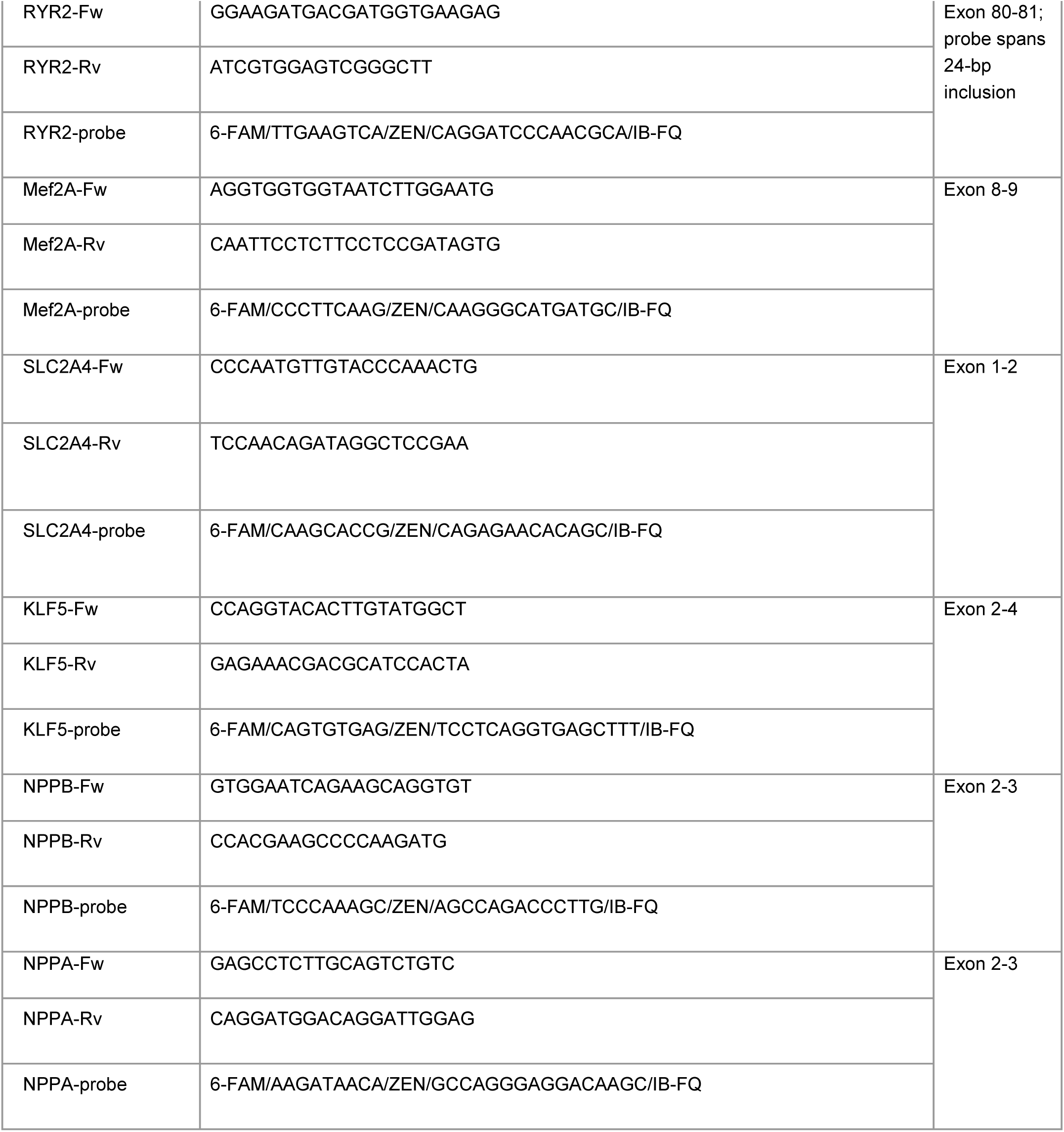
Oligos and primers used in this study.

**Table 2.** Plasmids used in this study.

| Name | Description |
| --- | --- |
| NG-ABE8e-BFP | A to G adenine base editor; NG-ABE8e (Addgene #138491) encoding ecTadA(8e)-nSpCas9(NG)-BFP |
| pBS | pBlueScript II SK(+); U6 promoter driving sgRNA expression with *Esp3I* cloning sites |
| pBS-ABE-sgRNA | pBS vector expressing the E913K-targeting ABE sgRNA |
| pCMV-PEmax-P2A-GFP | Mammalian expression vector for the SpCas9 PEmax prime editor fused to P2A-EGFP (Addgene #180020) |
| pU6-pegRNA-GG-acceptor | pegRNA acceptor plasmid with *Bsa*I sites (Addgene #132777) |
| PE-pegRNA | pegRNA construct generated by Golden Gate assembly following ref [2] Anzalone et al. (Nature, 2019) |
| pCMV-RBM20-tGFP | Wild-type RBM20 cDNA expressed under the CMV promoter with C-terminal turboGFP |
| pCMV-RBM20-E886G-tGFP | RBM20 E886G variant introduced into pCMV-RBM20-tGFP via site-directed mutagenesis (SDM) |
| pCMV-RBM20-W900R-tGFP | W900R variant introduced by SDM |
| pCMV-RBM20-P902R-tGFP | P902R variant introduced by SDM |
| pCMV-RBM20-N904Y-tGFP | N904Y variant introduced by SDM |
| pCMV-RBM20-M905I-tGFP | M905I variant introduced by SDM |
| pCMV-RBM20-E906V-tGFP | E906V variant introduced by SDM |
| pCMV-RBM20-V909L-tGFP | V909L variant introduced by SDM |
| pCMV-RBM20-E913K-tGFP | E913K variant introduced by SDM |
| pCMV-RBM20-E918Q-tGFP | E918Q variant introduced by SDM |
| pCMV-RBM20-E918D-tGFP | E918D variant introduced by SDM |
| pCMV-RBM20-E930K-tGFP | E930K variant introduced by SDM |
| pCMV-RBM20-D939N-tGFP | D939N variant introduced by SDM |

### Quantitative real-time PCR

Total RNA was extracted using the Ambion PARIS Kit (Invitrogen # AM1921). cDNA synthesis was performed with the SuperScript VILO cDNA Synthesis Kit (Invitrogen) using 2 µg of purified RNA as input. Gene expression was quantified by TaqMan real-time PCR using gene-specific primers and probes labeled with 6-FAM/ZEN/IBFQ (Table 1). Reactions were carried out with PrimeTime Gene Expression Master Mix (IDT) following the manufacturer’s instructions. Cycle threshold (Ct) values were obtained using the ViiA 7 Real-Time PCR System (Applied Biosystems), and relative mRNA expression levels were calculated using the 2^–ΔCt method.

### Western blotting

For detection of RBM20-tGFP fusion proteins in HeLa cells, HeLa cells were transfected with plasmids encoding different RBM20 variants using Lipofectamine 3000 (Invitrogen) in 6 well plates. For variant co-transfection experiments, a ratio of 1:1 was used per variant construct to add up to 500 ng total DNA per well. Twenty-four hours after transfection, cells were lysed in RIPA Lysis and Extraction Buffer (Pierce) and mixed with 2× Laemmli Sample Buffer (Bio-Rad) according to the manufacturers’ protocols. Protein samples were resolved on 4-20% Mini-PROTEAN TGX precast gel (Bio-Rad) and transferred to PVDF membranes using the Trans-Blot Turbo Transfer System (Bio-Rad). Membranes were probed with the following primary antibodies: mouse anti-turboGFP monoclonal antibody, clone OTI2H8 (OriGene, TA150041; 1:1,000), and mouse anti-β-actin monoclonal antibody, clone 937210 (BD, MAB8969; 1:10,000). IRDye 800CW anti-mouse secondary antibody (LI-COR, 925-32210; 1:10,000) was used for detection. Blots were visualized using the LI-COR Odyssey Fc imaging system and quantified with ImageJ.

### Immunocytochemistry (ICC)

Cells were seeded onto poly-D-lysine coated coverslips (Neuvitro, GG-12-1.5-PDL) in 12-well plates. For iPSC-derived cardiomyocytes, plates were additionally coated with Matrigel prior to cell seeding. Cells were fixed with 4% paraformaldehyde (PFA) and permeabilized with 0.2% Triton X-100 in PBS, followed by blocking with 10% goat serum in PBS-T (0.05% Triton X-100 in PBS). Coverslips were immunostained with a rabbit polyclonal anti-RBM20 primary antibody (Novus, NBP2-34038; 1:100 dilution) and a goat anti-rabbit IgG (H+L) Alexa Fluor Plus 488 secondary antibody (Thermo Fisher Scientific, A32731TR; 1:500). Coverslips were mounted using VECTASHIELD Vibrance Antifade Mounting Medium with DAPI (Vector Laboratories, H-1800). Images were acquired using a Leica SP8 inverted confocal microscope.

### Site-directed mutagenesis (SDM)

Customized mutagenesis services from GENEWIZ (Azenta) were used to introduce the exon 11 variants into the pCMV-RBM20-tGFP construct (Table 2).

### Cycloheximide (CHX) pulse-chase assay

CHX pulse-chase assays were used to determine the half-life of RBM20 variants in HEK293T cells. Equal amounts of pCMV-RBM20-tGFP plasmids encoding different RBM20 variants were transfected into HEK293T cells using Lipofectamine 3000 (Invitrogen). Cycloheximide (CHX, 200 µg/mL) was added at 0, 1, 3, 6, 10, and 24 hours post-transfection to inhibit protein synthesis. At each time point, cells were fixed with 4% PFA, stained with NucBlue Fixed Cell ReadyProbes Reagent (DAPI; Invitrogen), and transferred at 100 µL per well into black-wall, clear-bottom 96-well plates. Fluorescence was quantified using a SpectraMax iD3 plate reader with Ex/Em settings of 472/512 nm for tGFP and 360/460 nm for DAPI. tGFP fluorescence, reflecting RBM20 protein levels, was normalized to DAPI as a control for cell number, and protein half-life was determined by fitting normalized values to a one-phase exponential decay model with 95% confidence intervals.

### Hybrid enrichment of known RBM20 targets and RNA-sequencing for splicing

RNA was extracted from iPSC-CMs at 30-60 days of maturity using a mirVana isolation kit (Thermofisher). First strand synthesis was performed followed by hybrid capture with a custom panel (IDT, genes listed in Supplemental Figure 3B). RNAseq library was prepped and sequenced commercially using illumina products.

RMATS(26) was used for downstream analysis. Fastq files were quality-filtered and aligned to the human reference genome (GRCh38/hg38) using STAR v2.5.4b in two-pass mode with GENCODE v34 annotation and alignments were stored in BAM format. Alternative splicing events were detected by feeding BAM files into the software package rMATS, which, for every exon in the annotation set provided, calculates “percent spliced in” (PSI) values for each sample and outputs a ΔPSI value that quantifies the difference in the isoform ratio between two conditions. In this analysis, the alternative splicing events that were looked at were skipped exons (SE), retained introns (RI), and mutually exclusive exons (MXE). Alternative 5’ and 3’ splice site events were excluded.

To generate a deltaPSI score, which represents the difference in exon inclusion between wild and condition of interest, across genomic regions for specific genes of interest, we first average the predominant alternative splicing event, skipped exons, across biologic replicates to calculate mean exon inclusion percentages (defined as PSI within rMATS) normalized to read count. This PSI value can be matched with other samples by merging on identical genomic regions to subsequently generate deltaPSI values. deltaPSI values are then visualized by linking the corresponding genomic track and significantly spliced exons plotted using a modified autoplot function from the ggbio package (v1.45). Additionally, the absolute value of deltaPSI for each genomic region within a condition and its respective control are summed and divided by the total number of significant regions to generate an average deltaPSI per exon used to compare the average rate of differential splicing between conditions.

### AAV treatment and Kinetic Image Cytometry

AAV6-RBM20 (Supplemental Figure 3E), AAV6-GFP (Stanford Medicine’s Gene Vector and Virus Core (GVVC)) and AAV6-empty vector (***) were applied at indicated MOI at least 5 days after seeding of iPSC-CMs. Five days prior to AAV treatment, iPSC-CMs at least 40 days of differentiation using lactate media and metabolic maturation media(16) (beginning at day 20) were seeded to clear bottom optical 96 well plates. Five days after seeding, media was changed to that containing AAV (or control). Empty AAV was used as a control for calcium fluorescence experiments given interaction of AAV-GFP with the Cal-520 fluorophore. Five or 14 days following AAV treatment, cells were washed with 4x half media changes of FluoroBrite DMEM and a final x half media change with FluoroBrite DMEM, Cal-520 (AAT biosciences) and TMRM. The cells were then incubated for 1 hour at 37 °C and 5% CO_2_. After incubation, the cells were washed with 4 x half media changes of FluoroBrite DMEM and then returned to the incubator for 30 minutes. All media used was preheated in a water bath to 37 °C. After incubation, the cells were imaged on Kinetic Image Cytometer with interleaved acquisition of the movement of the cells measured in the TMRM channel (contractility) and fluorescence intensity of Cal-520 (calcium transients). Contractility and calcium measurements were extracted from the raw movies using in house Matlab scripts.

